# Genetic and behavioral adaptation of *Candida parapsilosis* to the microbiome of hospitalized infants revealed by *in situ* genomics, transcriptomics and proteomics

**DOI:** 10.1101/2020.03.23.004093

**Authors:** Patrick T. West, Samantha L. Peters, Matthew R. Olm, Feiqiao B. Yu, Yue Clare Lou, Brian A. Firek, Robyn Baker, Alexander D. Johnson, Michael J. Morowitz, Robert L. Hettich, Jillian F. Banfield

## Abstract

*Candida parapsilosis* is a common cause of invasive candidiasis, especially in newborn infants, and infections have been increasing over the past two decades. *C. parapsilosis* has been primarily studied in pure culture, leaving gaps in understanding of its function in microbiome context. Here, we reconstructed five unique *C. parapsilosis* genomes from premature infant fecal samples and analyzed their genome structure, population diversity and *in situ* activity relative to reference strains in pure culture. All five genomes contain hotspots of single nucleotide variants, some of which are shared by strains from multiple hospitals. A subset of environmental and hospital-derived genomes share variants within these hotspots suggesting derivation of that region from a common ancestor. Four of the newly reconstructed *C. parapsilosis* genomes have four to sixteen copies of the gene RTA3, which encodes a lipid translocase and is implicated in antifungal resistance, potentially indicating adaptation to hospital antifungal use. Time course metatranscriptomics and metaproteomics on fecal samples from a premature infant with a *C. parapsilosis* blood infection revealed highly variable *in situ* expression patterns that are distinct from those of similar strains in pure cultures. For example, biofilm formation genes were relatively less expressed *in situ*, whereas genes linked to oxygen utilization were more highly expressed, indicative of growth in a relatively aerobic environment. In gut microbiome samples, *C. parapsilosis* coexisted with *Enterococcus faecalis* that shifted in relative abundance over time, accompanied by changes in bacterial and fungal gene expression and proteome composition. The results reveal potentially medically relevant differences in Candida function in gut vs. laboratory environments, and constrain evolutionary processes that could contribute to hospital strain persistence and transfer into premature infant microbiomes.

## INTRODUCTION

Candida species are the most common cause of invasive fungal disease (Naglik et al. 2008; Silva et al. 2012). A variety of Candida species cause candidiasis and are recognized as a serious public health challenge, especially among immunocompromised and hospitalized patients (Clerihew et al. 2007, Bliss, 2015). Historically, *Candida albicans* most commonly has been recognized as the cause of candidiasis, and as a result, is the focus of the majority of Candida research (Kuhn et al. 2004; Trofa et al. 2008; Bliss, 2015). However, *Candida parapsilosis*, despite being considered less virulent than *C. albicans*, is the Candida species with the largest increase in incidence since 1990 (Trofa et al. 2008). Given important differences in the biology of *C. albicans* compared to non-albicans species, more research on non-albicans Candida species, especially the subset that poses a serious health risk, is needed (Bliss, 2015).

*C. parapsilosis* is often a commensal member of the gastrointestinal tract and skin (Trofa et al. 2008; Gonia et al. 2017). Passage from hospital workers’ hands to immunocompromised patients is thought to be a common cause of opportunistic infection in hospital settings (Huang et al. 1998). *C. parapsilosis* infections of premature infants are of particular concern. Indeed, *C. parapsilosis* is the most frequently isolated fungal organism in many neonatal intensive care units (NICUs) in the UK (Clerihew et al. 2007) and is responsible for up to one-third of neonatal Candida bloodstream infections in North America (Fridkin et al. 2006). Adding to the concern is the limited number of antifungal drugs and the increasing prevalence of antifungal drug resistance in Candida species. An estimated 3-5% of *C. parapsilosis* are resistant to fluconazole, the most commonly applied antifungal (Whaley et al. 2017). The recent emergence of multidrug-resistant *Candida auris* with its resultant high mortality rate (Forsberg et al. 2019) serves as a warning regarding the potential for outbreaks of multidrug-resistant *C. parapsilosis*. Therefore, understanding behavior of *C. parapsilosis*, both as a commensal organism and opportunistic pathogen, is incredibly important.

A challenge that complicates understanding of the medically relevant behavior of Candida in the human microbiome is that the hosts used in model infection systems (e.g., rat or murine mucosa) are not natural hosts to Candida species. Study of Candida in these models relies on some form of predisposition of the animal by occlusion, immunosuppression, surgical alteration, or elimination of competing microbial flora (Naglik et al. 2008). Pure culture experiments, an alternative to model system studies, are often the most accessible way to study Candida. However, the lack of a microbial community context is a large caveat, considering bacteria could influence the nutrition, metabolism, development, and evolution of eukaryotes. Indeed, other microbial eukaryotes have been shown to be dramatically influenced by their surrounding microbial communities. Choanoflagellates, the closest known living relative of animals, live in aquatic environments and feed on bacteria by trapping them in their apical collar (Hibberd et al. 1975). The Choanoflagellate S*alpingoeca rosetta* is primarily a unicellular organism but formation of multicellular rosettes is induced by a sulphonolipd (RIF1) and inhibited by a sulfonate-containing lipid, both produced by the bacterium *Algoriphagus machipongonensis* (Cantley et al. 2016). Furthermore, the bacterium *Vibrio fischeri* produces a chondroitinase, EroS, capable of inducing sexual reproduction in *S. rosetta* (Woznica et al. 2017). Together, these results demonstrate the influence that bacteria can exert on the morphology, development, and evolution of microbial eukaryotes.

There is more direct evidence motivating study of *C. parapsilosis* functioning *in situ*. For instance, *Caenorhabditis elegans* model of polymicrobial infection experiments showed that *C. albicans* exhibits complex interactions with *Enterococcus faecalis*, a bacterial human gut commensal and opportunistic pathogen. In this context, *C. albicans* and *E. faecalis* negatively impact one another’s virulence (Cruz et al. 2013), suggesting a mechanism that promotes commensal behavior in a gut microbial community context. The decrease in *C. albicans* virulence was attributed to inhibition of hyphal morphogenesis and biofilm formation by proteases secreted by *E. faecalis* (Cruz et al. 2013) as well as E. faecalis capsular polysaccharide (Bachtiar et al. 2016). No research has investigated *C. parapsilosis* in a microbial community context.

An alternative to studying Candida species in animal models or laboratory cultures is to use an untargeted shotgun sequencing approach (genome-resolved metagenomics). DNA is extracted from fecal or other samples and sequenced. The subsequent DNA sequences are assembled, and metagenome-assembled genomes (MAGs) are reconstructed. Much work of this type has focused on the bacterial members of the human microbiome; however, recently developed methods such as EukRep (West et al. 2018) enable reconstruction of eukaryotic genomes from metagenomes with greater consistency, including genomes of *Candida* species (Olm et al. 2019). The availability of genomes enables evolutionary studies and the application of other ‘omics’ approaches, such as transcriptomics, proteomics, and metabolomics, making it possible to go beyond metabolic potential to study activity *in situ*. Although there are limitations related to establishing causality via experimentation, the approaches can provide insights into metabolism and changes in metabolism linked to shifts in community composition in human-relevant settings.

Here, we applied shotgun metagenomics, metatranscriptomics, and metaproteomics to investigate the behavior and evolution of Candida in the premature infant gut and hospital environment. Novel *de novo* assembled *C. parapsilosis* and *C. albicans* genomes were reconstructed and the metagenomic data analyzed in terms of heterozygosity and population diversity. Due to the substantially less prior research on *C. parapsilosis* and the availability of *C. parapsilosis*-containing samples suitable for transcriptomics and proteomics, we focused our analyses on *C. parapsilosis* and identified genes and genomic regions under diversifying selection. Notably, we also identified instances of copy number gain of a gene involved in fluconazole resistance, pointing to a mechanism for hospital adaptation (Whaley et al. 2016). *C. parapsilosis in situ* transcriptomic and proteomic profiles were clearly distinct from profiles reported previously from culture settings. Substantial shifts in *C. parapsilosis* expression occurred with changes in microbiome composition over a few day period, suggesting the strong influence of bacterial community composition on *C. parapsilosis* behavior.

## RESULTS

### Recovery of novel Candida strain genomes

Fecal samples were collected from 161 premature infants primarily during the first 30 days of life (DOL) (full range of DOL 5–121), with an average of 7 samples per infant. Samples of the Neonatal Intensive Care Unit (NICU) were taken from six patient rooms within the hospital housing the infants (Magee-Womens Hospital of UPMC, Pittsburgh, PA, USA). Candida genomes were assembled from samples containing >2 Mbp of predicted eukaryotic DNA using a EukRep-based pipeline (West et al. 2018; see the “Methods” section for details). Three of the Candida genomes (Olm et al. 2019) and the bacterial component (Olm et al. 2019) were analyzed previously (see the “Methods” section). Eleven unique Candida genomes were assembled in total (Table 1), six *C. albicans* genomes and five *C. parapsilosis* genomes. All genomes have an estimated completeness >85% except for *C. parapsilosis* L2_023 and NYC subway, which had low coverage (4x and 6x respectively) in their samples. Nine genomes were reconstructed from premature infant fecal samples; one genome was derived from a NICU room sample S2_005. For comparison, we analyzed a Candida genome that we reconstructed from a publically available metagenome read dataset from the New York City subway (NYC_subway; Afshinnekoo et al. 2015), as well as four previously published *C. parapsilosis* and fifty-one *C. albicans* isolate genomes.

**Table 1:** Overview of Candida strain genomes used in this study.

| Genome | Genus | Species | Length | # Scaffolds | N50 | BUSCO Comp. | Sample type | Reference |
| --- | --- | --- | --- | --- | --- | --- | --- | --- |
| C1_006 | Candida | parapsilosis | 11852211 | 191 | 108686 | 92 | Infant fecal metagenome | This study |
| N3_182 | Candida | parapsilosis | 12563647 | 342 | 65710 | 94 | Infant fecal metagenome | Olm et al. 2019 |
| S2_005 | Candida | parapsilosis | 11573959 | 1051 | 14507 | 93 | NICU metagenome | Olm et al. 2019 |
| NYC Subway | Candida | parapsilosis | 7420453 | 1285 | 6417 | 62 | NYC subway metagenome | This study |
| L2_023 | Candida | parapsilosis | 4870205 | 2906 | 1700 | 35 | Infant fecal metagenome | This study |
| CDC317 | Candida | parapsilosis | 13030174 | 9 | 2091826 | 93 | Clinical skin isolate | Butler et al. 2009 |
| GA1 | Candida | parapsilosis | 13025060 | 39 | 1114083 | 93 | Clinical human blood isolate | Pyrzycz et al. 2013 |
| CBS1984 | Candida | parapsilosis | 13044404 | 25 | 962200 | 92 | Olive fruit isolate | Pyrzycz et al. 2013 |
| CBS6318 | Candida | parapsilosis | 13050515 | 28 | 1691491 | 93 | Healthy skin isolate | Pyrzycz et al. 2013 |
| N1_023 | Candida | albicans | 13456346 | 1675 | 15180 | 94 | Infant fecal metagenome | This study |
| N2_070 | Candida | albicans | 13540857 | 1614 | 14761 | 93 | Infant fecal metagenome | This study |
| N5_264 | Candida | albicans | 11647081 | 746 | 27434 | 85 | Infant fecal metagenome | This study |
| S3_003 | Candida | albicans | 11972257 | 1049 | 14710 | 87 | Infant mouth metagenome | This study |
| S3_016 | Candida | albicans | 10068784 | 802 | 19749 | 86 | Infant mouth, skin, and gut metagenome coassembly | This study |
| SP_CRL | Candida | albicans | 12561678 | 897 | 22840 | 91 | Infant fecal metagenome | Olm et al. 2019 |

### Candida genomic variability

To characterize genomic variability in the strains of *C. albicans* and *C. parapsilosis* represented by metagenome-derived genomes, we identified single-nucleotide variants (SNVs) by mapping reads against completed reference genomes (strain SC5314 for *C. albicans* and CDC317 for *C. parapsilosis*). *C. albicans* genomes ranged from 3.2-9.9 heterozygous SNVs per kb (heterozygosity), whereas *C. parapsilosis* genomes ranged from 0.12-0.38 heterozygous SNVs per kb. Thus, we infer that, compared to *C. albicans, C. parapsilosis* displays very low variability in its diploid chromosome pair, which can be indicative of low genetic variability in the hospital environment and primarily asexual reproduction (Magwene et al. 2011).

Low heterozygosity in *C. parapsilosis* genomes has been reported for previously sequenced genomes (Pryszcz et al. 2013). Interestingly, *C. parapsilosis* genomes derived from our fecal metagenomes showed even lower overall heterozygosity than pure culture reference genomes (Figure S1). In general, this would not be expected because within-sample population diversity due to sampling of a microbial community should inflate measures of genomic heterozygosity. Thus, the lower genomic heterozygosity may be reflective of infants being initially colonized by essentially a single Candida genotype.

Because multiple new strains were sequenced from the same hospital, the phylogenetic relationships of new and previously sequenced strains from the same hospital were of interest from the perspectives of the persistence of Candida populations in the hospital environment and transfer from room to human. To place the hospital and gut-associated sequences in context, we first compared those genomes to available reference genomes from NCBI using pair-wise average nucleotide identity (ANI) and by construction of single nucleotide variant (SNV) trees (Figure 1A, Figure S1-2). L2_023 was not included due to low sequencing coverage. *C. albicans* strains were spread throughout the tree of known *C. albicans* diversity (Figure S2) whereas *C. parapsilosis* strains from infant gut and NICU samples were clustered on a single branch (Figure 1A) separate from other reference hospital and environmental strains. Further, the two infant gut strains, sampled years apart, were nearly identical (99.99% identity). We verified this with whole genome alignments of the hospital and gut sequences (Figure S1-S2). We thus infer that the hospital room and gut *C. parapsilosis* strains are very closely related.

**Figure 1:**
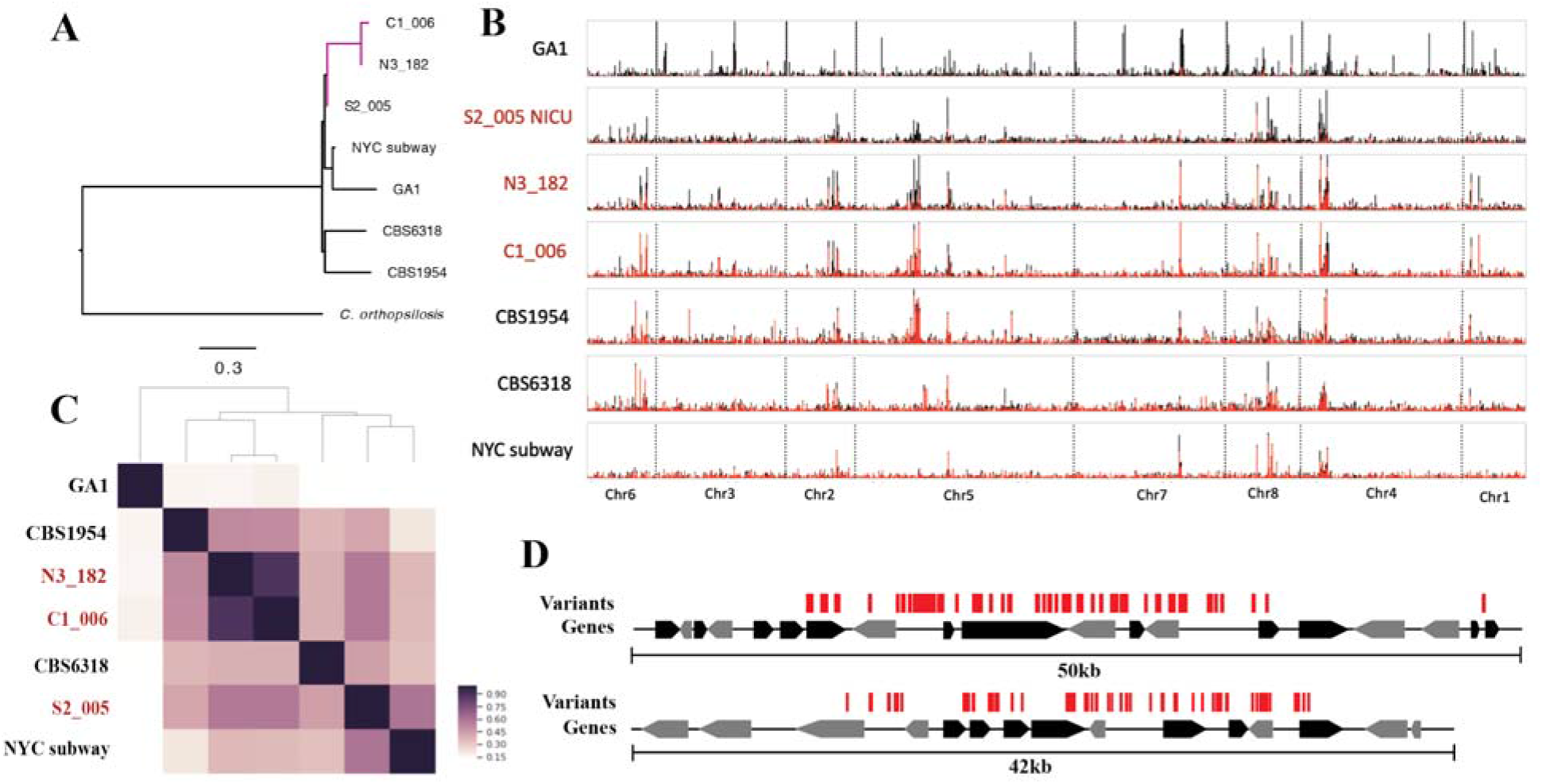
Analysis of *C. parapsilosis* genomic variability reveals a potential hospital associated population and the presence of SNV hotspots. (A) A phylogenetic tree of *C. parapsilosis* strains constructed from concatenated SNVs. Metagenome derived hospital strains from this study demarcated as the purple clade. ANI comparisons and a *C. albicans* SNV tree are also available in Figures S1-S2. (B) Whole genome SNV density plots for each *C. parapsilosis* strain. Strain names in red are strains assembled from samples from infants or the NICU from Magee-Women’s Hospital. SNV density plotted in 1.3kb sliding windows. Window size was selected based on ease of visualization. Chromosomes are separated with dashed lines. Total bar height represents total SNV density and homozygous SNV proportion is labeled in red whereas heterozygous is black. (C) Depiction of SNV hotspot overlap between each strain. Pairwise overlap was calculated between each strain and plotted. Strain names in red are strains assembled from samples from infants or the NICU from Magee-Women’s Hospital. (D) Two example SNV hotspots. Individual SNVs are represented with red bars.

Based on analysis of population structure of seven *C. parapsilosis* genomes (Figure S3), we predicted six distinct *C. parapsilosis* ancestral populations. The exception is the fecal strain N3_182, which appears to be a recombinant admixture of the ancestral populations NICU strain S2_005 and the fecal strain C1_006. Given that N3_182 was sequenced four years before C1_006, both parental strains must have both existed in the hospital environment prior to hybridization. The findings provide evidence for a clearly defined, distinct hospital associated *C. parapsilosis* strains, a hybrid of which colonized a premature infant.

### C. parapsilosis SNV hotspots as indicators of genes under selection

To investigate whether genomes sampled from the hospital could provide evidence of evolutionary adaptation to this environment, we visualized the spatial distribution of *C. parapsilosis* genomic diversity in the newly reconstructed genomes by mapping reads from each genome to a reference sequence (CDC317, recovered from a clinical sample) and calling SNVs. We plotted the density of SNVs in 1.3 kbp sliding windows across the genome of each strain (Figure 1B). Both heterozygous and homozygous SNVs are largely evenly distributed throughout the genome, with the exception of a few small regions with highly elevated SNV counts (regions of elevated diversity) that we refer to as SNV hotspots (Figure 1B).

Interestingly, SNV hotspots show a high level of conservation between all strains (Figure 1C). The one exception is reference strain GA1 cultured from human blood (Pryszcz et al. 2013), which shares only ~10% of its SNV hotspots with any other given strain. Notably, the NY subway strain is fairly similar to the clinical reference strain (few and minor hotspots) whereas our hospital sequences share SNV hotspots with both of the CBS strains (one from an olive and the other from skin), consistent with genomic similarity of the hospital and CBS strains in those regions.

To provide a more complete view of variation hotspots, we also mapped the reads from each population to the three other reference genomes (environmental strains CBS1984 and CBS6318, and the GA1 blood isolate, Figure S4). The number of SNV hotspots ranged from 16-45, and the regions were 5 kb to 24.5 kb in length. Due to the large size of the SNV hotspots, each hotspot overlaps a number of individual genes with SNVs spread both within and between genes (Figure 1D). In total, 376 genes are present within a SNV hotspot in at least one strain. No particular KEGG family or PFAM domain was significantly enriched in SNV hotspots. This, combined with the fact that SNVs are spread both within and between genes may be indicative of SNV hotspots being recombination hotspots, or locations where additional SNVs hitchhike along with SNVs under selection.

### Multicopy RTA3 gene

Another explanation for SNV hotspots could be due to gene copy number variation, as recent duplications of a region acquire mutations yet reads from these duplications map back to a single location. Overall, when windowed genomic coverage is plotted alongside SNV density (Figure 2A), this is clearly not the case. However, across the entire genome two regions of high coverage (Figure 2A), indicating high copy number variation, were identified and neither correspond to SNV hotspots. The first high copy number region contains an estimated 17-28 copies of the 18S, 25S, 5S, and 5.5S rRNA genes (Table S1, Figure 2B). The variation in rRNA copy number may indicate a range of maximum growth rates (Roller et al. 2016). The second region, which corresponds to the lipid translocase RTA3 gene and flanking sequence, is present in 5-16 copies (Table S1) in strains C1_006, N3_182, L2_023, S2_005, and NYC_subway but not the four reference genomes (Figure 2B). The high copy number RTA3 genes also have no detectable SNVs and different boundaries in each strain, suggesting the duplications were very recent and independent events in each strain.

**Figure 2:**
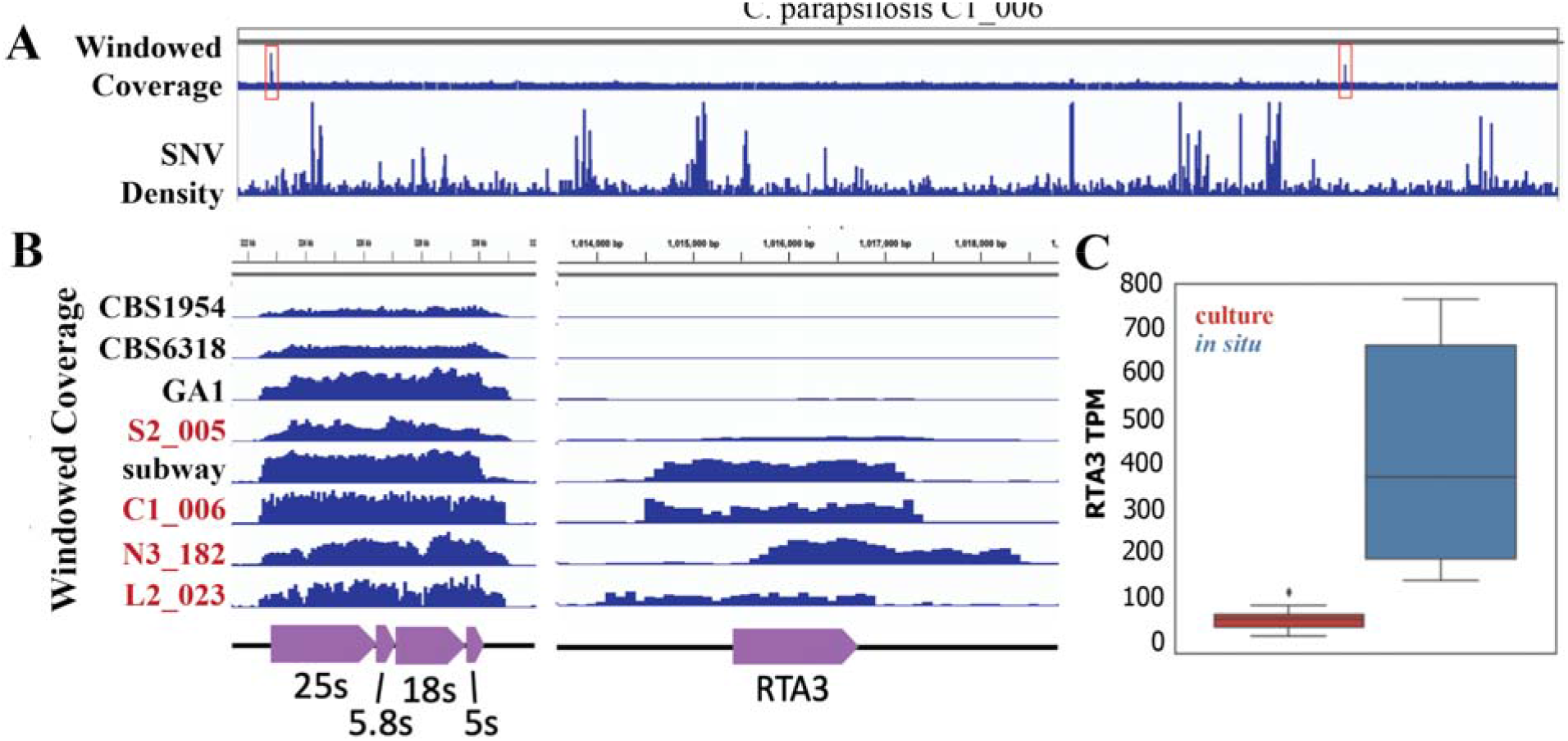
*C. parapsilosis* strains have high copy number rRNA and RTA3 loci. (A) Whole genome windowed coverage of SNP density for *C. parapsilosis* strain C1_006. High copy number regions of interest are highlighted with red boxes. (B) An expanded view of highlighted high copy number regions from (A). Windowed coverage is plotted as 100bp sliding windows. Metagenome-derived hospital strains from this study labeled in red. (C) Boxplot of expression of the RTA3 gene from multicopy strain C1_006 *in situ* (red) and strain CDC317 in culture (blue). Expression represented as Transcripts Per Million (TPM).

### In situ metatranscriptomics and metaproteomics

Given most work with Candida species is performed in pure culture or in murine models, little is known about their behavior in the human gut. We hypothesized performing metatranscriptomics and metaproteomics on infant fecal samples with *C. parapsilosis* would reveal unique transcriptomic and proteomic profiles, indicative of differences in metabolism and behavior between culture and *in situ* settings. Two prospective infants were identified, infant 06 with a documented Candida blood infection (Figure 3) and infant 74 with a documented Candida lung infection. Both infants were treated with fluconazole shortly after detection of Candida infection (Figure 3, Table S2). Metagenomic, metatranscriptomic, and metaproteomic datasets were generated from fecal samples at five to six timepoints for each infant. In infant 74, no Candida species were detected in the generated datasets (Figure S4). However, in infant 06, metagenomic sequencing confirmed the presence of *C. parapsilosis* (strain C1_006) in the fecal samples. *De novo* gene prediction was performed on the metagenome-derived *C. parapsilosis* genome and the resulting gene models were used for mapping transcriptomic reads and proteomic peptides (Figure 3).

**Figure 3:**
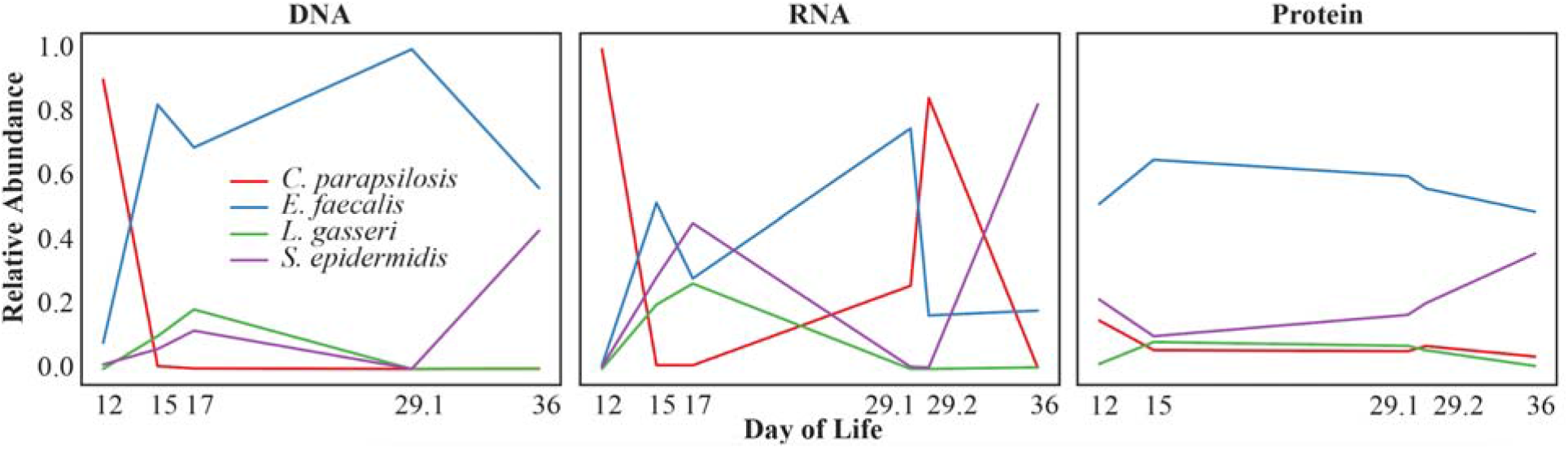
*In situ* metagenomics metatranscriptomics, and metaproteomics of infant 06. Plotted are the relative DNA, RNA, and peptide abundances for each detected organism after human removal. Plotted on the x axis are the Days Of Life (DOL) samples were taken.

In addition to *C. parapsilosis*, genomes were recovered for three bacterial species in infant 06: *Enterococcus faecalis*, *Lactobacillus gasseri*, and *Staphylococcus epidermidis*. Interestingly, in every infant where a Candida genome was assembled or detected through read mapping, *E. faecalis* was also present (N=7). *C. parapsilosis* is highly abundant in the first 20 days of life before quickly being replaced or outnumbered, largely by *E. faecalis*. Similar abundance patterns have been observed previously for microbial eukaryotes in neonatal fecal samples (Olm et al. 2019). *C. parapsilosis* transcriptomic abundance shows a similar pattern to the DNA abundance but transcription remains detectable at later time points (Figure 3). In contrast, *C. parapsilosis* proteomic abundance remained relatively stable over all timepoints.

### C. parapsilosis expression in situ vs. culture settings

Given most work with *C. parapsilosis* has been performed on pure cultures, we wondered if there are differences in behavior and metabolism that would be detectable by comparing transcriptomic datasets. For comparison, we downloaded raw sequencing reads from publically available *C. parapsilosis* RNAseq experiments (Guida et al. 2011; Pryszcz et al. 2013), including datasets from multiple strains (CDC317, CBS1954, and CBS6318) and varying culture conditions, including different media, growth temperatures, and oxygen concentrations. A hierarchical clustering of expression of CDC317 transcripts reveals a clearly distinct transcriptomic profile between *in situ* and all culture settings (Figure 4A). Notably, *in situ* samples are also extremely variable; clustering as far apart from one another as from the culture samples (FIgure 4A). We quantitatively identified differentially expressed transcripts between culture and *in situ* settings and found that 53% of transcripts were significantly differentially expressed; 23% up *in situ*, 30% down (Figure 4B), further highlighting the stark differences between *in situ* and culture settings.

**Figure 4:**
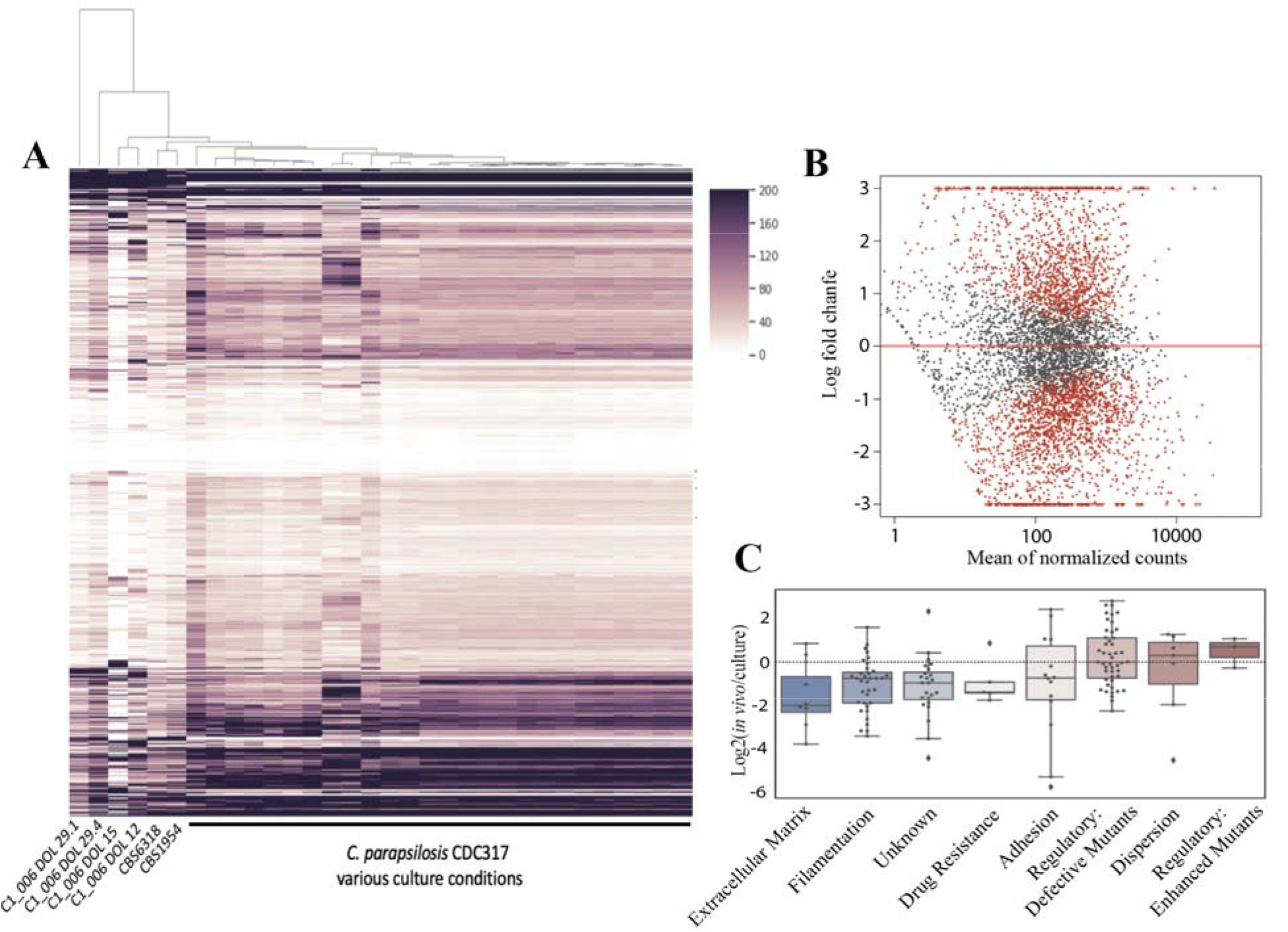
*C. parapsilosis* displays distinct and highly variable *in situ* transcriptomic profiles. (A) Hierarchical clustering of *C. parapsilosis* TPM values for C1_006 in *in situ* samples and pure culture samples under a variety of conditions. (B) Average log2 fold change *in situ* vs culture plotted against the mean of normalized counts for eac transcript. Transcripts in red were identified as being significantly differentially expressed by DESeq2. (C) Boxplots of expression of categories of genes involved in biofilm formation. Regulatory defective mutants refers to regulatory genes that inhibited biofilm formation when mutated.

*In situ* and culture transcriptome samples were differentiable in a principal component analysis (PCA), paralleling the hierarchical clustering of *C. parapsilosis* transcriptomes (Figure 5A). We performed a sparse Partial Least Squares Discriminant Analysis (sPLS-DA), treating each transcript as a variable, to try and identify important features able to discriminate between *in situ* and culture in a multivariate space (Figure 5B, Figure S5, Table S3). Important features were enriched for mitochondrial and aerobic respiration genes (9/50), uncharacterized genes (15/50), and a subset of ribosomal proteins (8/50; p=2.3×10^−7^).

**Figure 5:**
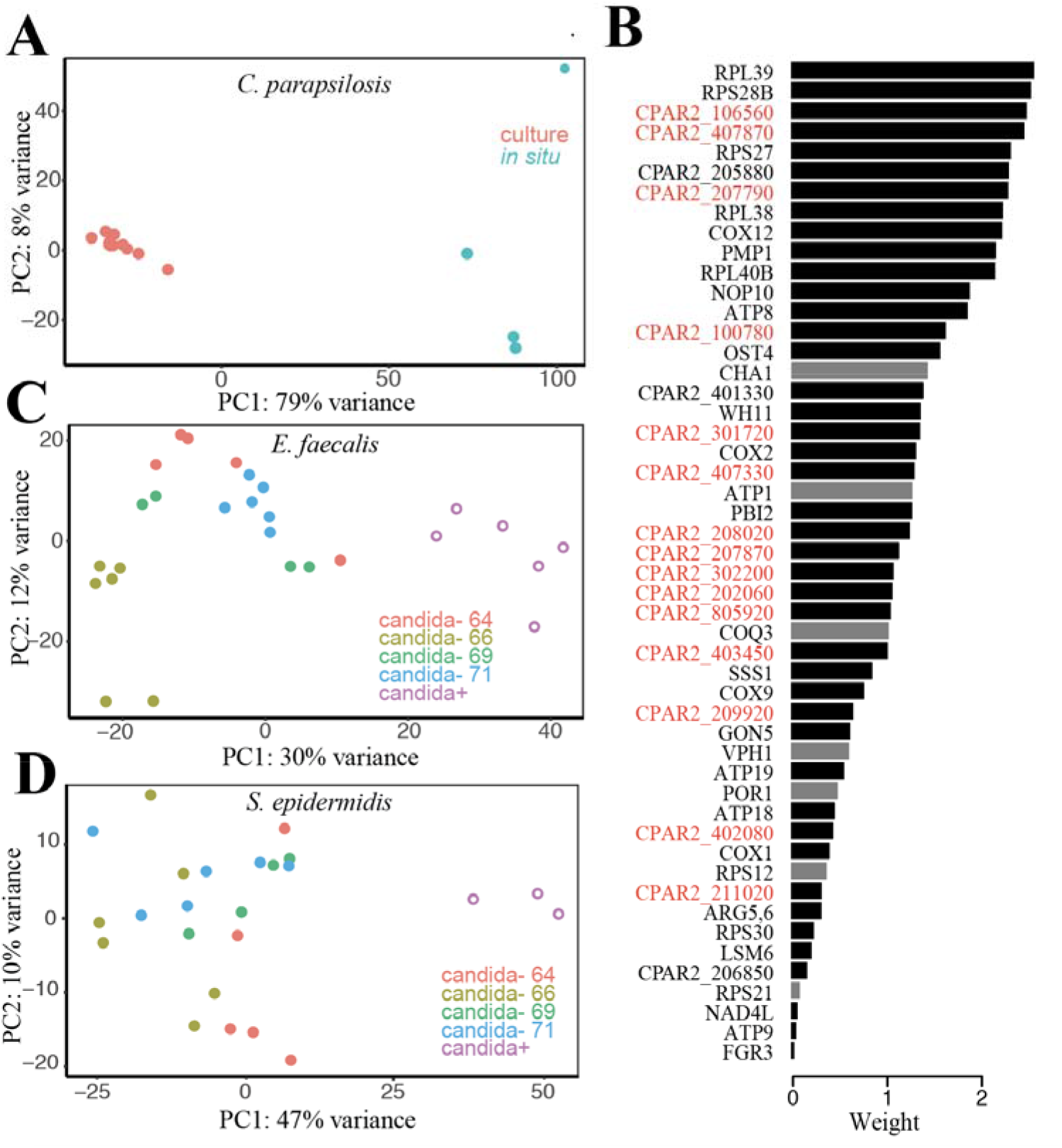
Presence of *C. parapsilosis* affects bacterial community member’s expression. (A) PCA of *C. parapsilosis in situ* and pure culture transcriptomes. (B) Depiction of features identified by sPLS-DA for separatin *C. parapsilosis in situ* and pure culture transcriptomes. Plotted are the feature weights. Black bars are genes that exhibited higher expression on average *in situ* whereas grey had higher average expression in culture. Genes labeled in red correspond to proteins of unknown function. (C-D) PCAs of *E faecalis* (C) and *S. epidermidis* (D) transcriptomes from infant microbiomes both with and without detected *C. parapsilosis*. Candida-negativ transcriptomes were from four different infants (published previously; Sher et al. 2020) denoted as 64, 66, 69, and 71.

Biofilm formation is an important virulence factor for Candida species; often contributing to the development of systemic infections (Nobile et al. 2012;Nobile et al. 2015). We were interested in whether the expression of virulence factors was enriched *in situ,* given the samples were obtained from an infant with a documented Candida blood infection. We obtained a list of well characterized biofilm formation genes from *C. albicans* (Nobile et al. 2015), identified orthologs in *C. parapsilosis* and compared their expression *in situ* to culture settings. Biofilm formation showed lower expression overall *in situ* (Figure 4C).

We were curious to see if the multicopy RTA3 gene in infant strain C1_006 (Figure 2B) showed increased expression as compared to the single copy RTA3 gene in reference strain CDC317. Indeed, the expression of the RTA3 in strain C1_006 is significantly higher (Figure 2C), suggesting a role of this gene duplication as a way to increase overall expression of RTA3. Interestingly, we did not see an increase in expression following fluconazole treatment (Figure S6), indicating RTA3 expression may be consistently higher in C1_006. However, it is worth noting we were unable to obtain samples until seven days after fluconazole treatment and any treatment effect on expression may have already passed.

### C. parapsilosis impact on bacterial expression

*E. faecalis*, *S. epidermidis* and *L. gasseri* bacteria in infant 06 had transcripts sequenced at high depths at multiple time points (Figure 3) so it was possible to investigate whether the presence or absence of *C. parapsilosis* had a distinguishable effect on their transcriptomic profiles. We compared bacterial transcription in these samples to transcription patterns of bacteria in the absence of Candida using previously reported datasets (21 samples for *E. faecalis* and 20 samples for *S. epidermidis;* Sher et al. 2020). The analysis was not possible for *L. gasseri* as this bacterium was not present in any of the metatranscriptomes used for comparison. The transcriptomes of both *E. faecalis* and *S. epidermidis* were distinguishable between the presence and absence of *C. parapsilosis*, and this effect appears to be independent of infant of origin and thus the bacterial strain variant type (Figure 5C-D). This result suggests *C. parapsilosis* has a large impact on the behavior and metabolism of other gut community members. In addition, the expression of *E. faecalis* genes previously shown to negatively impact *C. albicans* virulence (Cruz et al. 2013) showed no significant difference in expression between *C. parapsilosi*s negative and positive samples.

Important features identified from a sPLS-DA on Candida-positive vs. Candida-negative samples included a subset of *E. faecalis* ribosomal proteins (Table S3, Figure S5). Additionally, ribosomal proteins all showed higher expression *in situ*, suggesting increased *E. faecalis* growth rate in the presence of *C. parapsilosis*. Other important features included mannitol specific phosphotransferase system (PTS) transporters upregulated in Candida-positive samples and downregulated mannose specific PTS transporters (Table S3). Furthermore, Mannitol-1-phosphate 5-dehydrogenase, an enzyme responsible for the conversion of D-mannitol to fructose, was upregulated in Candida-positive samples, indicating an increased capacity for degradation of mannitol in addition to import (Table S3). Important features in *S. epidermidis* were less clear, but again included a subset of ribosomal proteins as well as beta-lactamases, both with increased expression *in situ* (Table S3).

### Transcriptomics enriched gene functions

Given the large differences in transcriptomes between culture and *in situ*, we looked for functions enriched in either setting (Figure 6, Table S4). DESeq2 identified groups of differentially expressed genes that were too large to be informative, so more restrictive cutoffs were used. Up *in situ* was defined as having >3 log2 expression *in situ* whereas down *in situ* was defined as <-3 log2 expression *in situ*. Up *in situ* was enriched for KEGG families for LSM 2-8 and 1-7 complexes, a family of proteins involved in mRNA metabolism highly conserved in eukaryotes (Beggs et al. 2005), as well as Cytochrome c oxidase and bc1 complex and proteins without an annotated KEGG family (Figure 6, Table S4). Down *in situ* is enriched for helicase and polysaccharide synthase PFAM domains. Additionally, proteins without an annotated KEGG family (unknown function) were enriched in both groups (Table S4).

**Figure 6:**
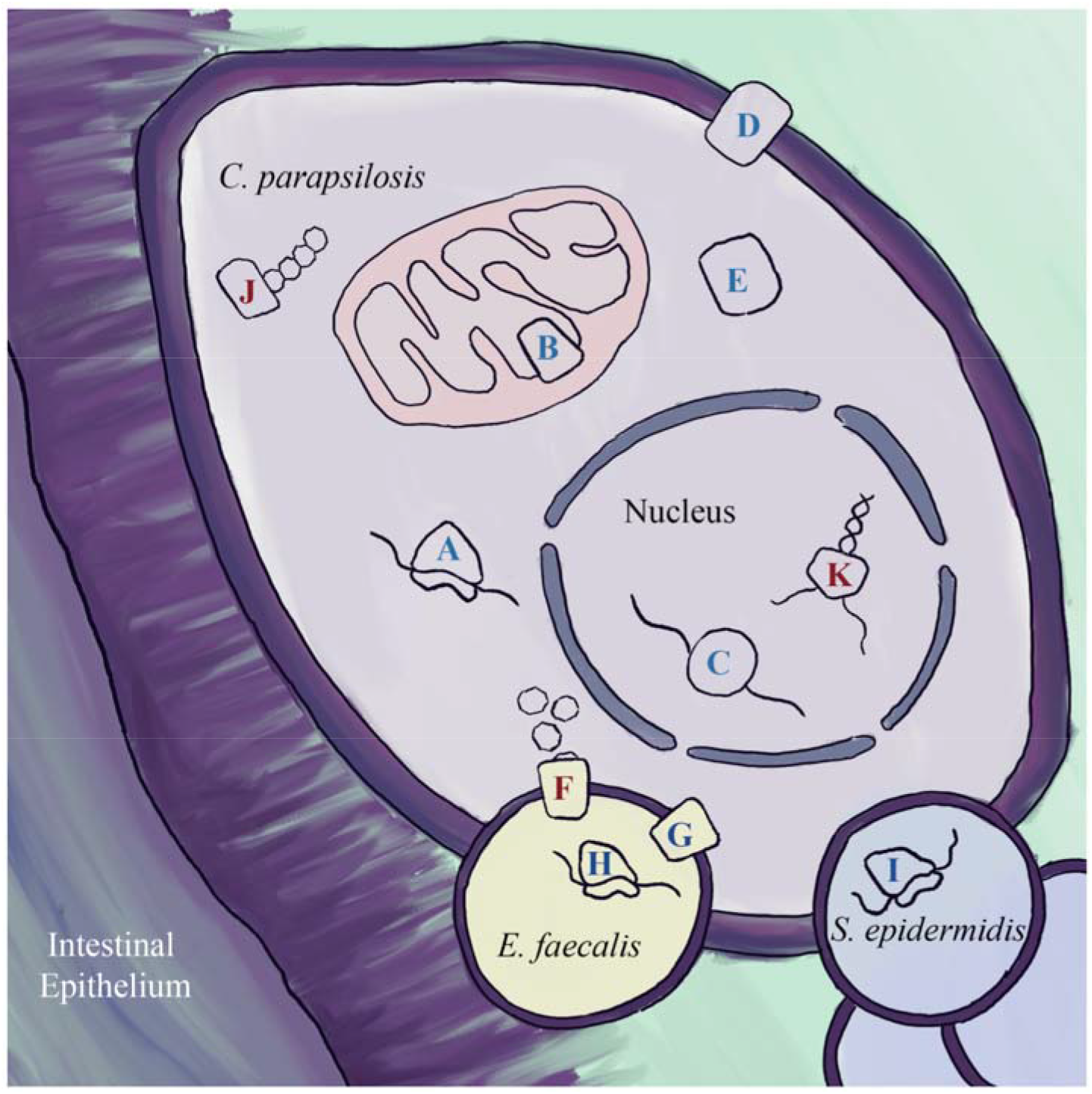
*in situ* enriched gene categories. Diagram depicting *C. parapsilosis* in the context of the infant gut, highlighting gene categories or families that were significantly enriched in differentially expressed genes between *in situ* and culture. Blue letters represent functions with higher expression *in situ*, while red represent functions wit lower expression *in situ*. See Table S5 for details. (A) ribosomal proteins (B) cytochrome c oxidase subunits (C) LSM complexes (D) proton antiporters (F) *E. faecalis* mannose transporters (G) *E. faecalis* mannitol transporters (H) *E. faecalis* subset of ribosomal proteins (I) *S. epidermidis* subset of ribosomal proteins (J) *C. parapsilosis* polysaccharide synthases (downregulated in situ) (K) *C. parapsilosis* helicases (downregulated *in situ*).

### Proteomics

As noted above, the metaproteomic abundances for infant 06 were relatively stable over time, with evidence of Candida core metabolic activities such as glycolysis, which indicate stability of this organism within the gut environment (Figure 3C). Comparison of the Candida *in situ* proteomics data with data from pure culture experiments was not possible as no pure culture proteomics datasets suitable for comparison have been published. Using the metagenome-derived *C. parapsilosis* genome as reference, we identified the most abundant proteins and found that this subset included ribosomal and F-Type ATPase proteins (Figure 6) and was significantly enriched for HSP70 and actin PFAM domains (Table S4). Also among proteins found with the most peptide evidence were proteins related to protection of the organism from oxidative stress, such as superoxide dismutase. The high abundance protein set included some of the genes contained within SNV hotspots but there was no significant association. We also examined the most abundant proteins in the bacterial species. In *E. faecalis* and *S. epidermidis*, Lac genes were some of the most abundant proteins suggesting lactose may be an important substrate for these community members. Finally, among human proteins detected, there was ample evidence of neutrophil degranulation, which indicates an active host immune response. Neutrophils use oxidative mechanisms to promote fungal clearance (Desai et al., 2018), which suggests Candida is employing oxidative protection in response to this host defense mechanism.

## DISCUSSION

Fungal pathogens are known to have hospital reservoirs. For example, the water supply system of a paediatric institute was shown to be a reservoir for *Fusarium solani* (Mesquita-Rocha et al., 2013). A NICU outbreak of the fungi *Malassezia pachydermatis* was linked to the dog of a healthcare worker (Chang et al., 1998), although persistence via long-term carriage by a healthcare worker vs. continual passage between infants and rooms (or a combination of these) could not be resolved. However, much remains to be learned about where reservoirs of hospital-associated fungi are and how long strains persist in them. In contrast to previous studies of *C. parapsilosis* utilizing pure culture and model systems, we applied genome-resolved metagenomics, metatranscriptomics, and metaproteomics to study *C. parapsilosis* in the context of the infant gut and hospital rooms of a neonatal intensive care unit. We detected novel, near identical *C. parapsilosis* genomes sequenced years apart in separate infants, suggesting transmission of members of a fungal population from reservoir to infant or infant to reservoir to infant. It is worth noting that although the strains are near-identical, the multicopy RTA3 locus in each strain had different boundaries and different copy numbers. This observation suggests that these two strains are very closely related members of a somewhat more diverse hospital adapted population.

Population genomic analyses of reconstructed genomes revealed multiple, independent instances of copy number gain of the RTA3 gene. RTA3, a lipid translocase, has been implicated in resistance to azole class antifungal drugs such as fluconazole in *C. albicans* (Whaley et al. 2016). The RTA3 gene is frequently overexpressed in resistant isolates and increased expression of RTA3 increases resistance to fluconazole whereas deletion of the RTA3 gene results in increased azole susceptibility (Whaley et al. 2016). Copy gain of this gene in *C. parapsilosis* strains may represent a mechanism for rapid adaptation to fluconazole, the most widely used antifungal in most hospitals (Whaley et al. 2016), as a means by which to increase its expression and thus its resistance. Similar gene copy number gains have been reported for the human amylase gene, hypothesized to be in response to increases in starch consumption (Pajic et al. 2019). Indeed RTA3 expression *in situ* from strain C1_006, which has RTA3 in multicopy, was significantly increased as compared to single copy strain CDC317 in culture (Guida et al. 2011; Figure 2C). The high likelihood that the copy number gain occurred independently in multiple strains suggests selection for this particular genomic feature. Identifying mechanisms of antifungal resistance is of particular importance given 3-5% of *C. parapsilosis* strains are already resistant to fluconazole (Whaley et al. 2017) and our relative inability to deal with infections of drug-resistant fungi.

Examining the genomic distribution of SNVs within the genomes of each *C. parapsilosis* strain revealed the presence of SNV hotspots. Many of these SNV hotspots are shared between strains, some of which are specific to the hospital and infant gut strains. Unlike *C. albicans*, *C. parapsilosis* is not an obligate commensal of mammals (Trofa et al. 2008). Consequently, some regions of the *C. parapsilosis* genome may be under selection for adaptation to the hospital, in addition to the gut environment. Further supporting the idea that some genomic innovation is associated with adaptation to the built environment, the NICU strain clustered the most closely to the NYC subway strain based on SNV hotspot overlap (Figure 1C). These two strains are geographically and phylogenetically distinct but the shared regions of diversification may be related to their common need to adapt to the built environment.

Metatranscriptomics of infant fecal samples revealed *C. parapsilosis* transcriptomes that are both highly variable and distinct from those of culture samples. Interestingly, the degree of variance exhibited by transcriptomes of the same population in the same infant over a few day period was greater than that observed between *C. albicans* white and opaque phenotypes (Figure S7; Tuch et al. 2010). The *C. albicans* white and opaque phenotypes differ in their appearance (Slutsky et al. 1987), mating style (Miller et al. 2002), and environmental conditions they are best adapted to (Ramirez-Zavala et al. 2008, Huang et al. 2009), and represent two exceptionally distinct Candida phenotypes. The high variability in *C. parapsilosis* is likely the result of changing conditions presented in the gut, including microbial community composition as well as the developing physiology of the host. Varying stages of infection and/or response to antifungal treatment may also have had an effect, but more dense time-series and additional infants would be required to elucidate these effects.

In contrast to the large changes in *C. parapsilosis* RNA and DNA relative abundances over time, *C. parapsilosis* peptide relative abundance remained stable over the study period. It is not uncommon to see different signals from transcripts and proteins (Haider et al. 2013), in part because proteins can persist for relatively long periods of time compared to transcripts. The most abundant proteins in the proteomics dataset have a HSP70 domain found in heat shock proteins (HSP). In *C. albicans*, HSP have been documented to help control virulence by interacting with regulatory systems, and to enable drug resistance (Gong et al. 2017).

The presence of *C. parapsilosis* within infant gut samples may impact the transcriptomes of bacterial gut community members. Important features for separating Candida-positive and Candida-negative samples included a suite of upregulated mannitol transporters and downregulated mannose transporters in *E. faecalis* (Table S3). *C. parapsilosis* strain SK26.001 is documented as producing mannitol (Meng et al. 2017) and mannose, in the form of the polysaccharide mannan, which can be an important component of extracellular polysaccharides produced by Candida (Dominguez et al. 2019). Interestingly, a characteristic of *E. faecalis* is its ability to grow by fermenting mannitol (Quiloan et al. 2012). Given the potential for interaction between *E. faecalis* and *C. parapsilosis*, its possible the disappearance of *C. parapsilosis* induced a substrate switch in *E. faecalis*.

Interestingly, statistical tests detected a subset of ribosomal proteins as important features for separating transcriptome patterns of *C. parapsilosis in situ* from those reported from culture studies, as well as for separating Candida-positive from Candida-negative samples for both *E. faecalis* and *S. epidermidis* (Table S3). In recent years, ribosomal heterogeneity, in which ribosomal protein subunits are swapped out or missing from individual ribosomes, has gained traction as a way for organisms to regulate translation (Guimaraes et al. 2016, Shi et al., 2017, Genuth et al. 2018). Ribosomal heterogeneity may be being utilized as an additional regulatory measure to adapt to the rapidly changing gut microbial context. Alternatively, fluctuations in ribosomal subunit abundance could be to maintain ribosomal homeostasis (Cruz et al. 2017), or individual ribosomal subunits could be performing functions unrelated to protein synthesis (Zhou et al. 2015).

Biofilm formation is an important virulence factor of Candida infections (Cavalheiro et al. 2018). Infant 06 had a documented Candida blood infection, and such infections are commonly systemic (Mavor et al. 2005). Interestingly, despite infection, Candida biofilm formation genes were relatively less expressed *in situ* in the gut of Infant 06 as compared to expression levels previously reported over a range of culture conditions. Similarly, genes with a PFAM domain for polysaccharide synthase, genes potentially important for the generation of Candida biofilm matrices (Dominguez et al. 2019), were less expressed *in situ* than in cultures. Thus, biofilm formation may not be an important component of every infection.

Genes linked to oxygen utilization, such as cytochrome c oxidase subunits, were more highly expressed *in situ* than over the range of culture conditions, suggesting growth in a relatively aerobic environment. This may be reflective of the higher oxygen levels in the gut during early life (Chong et al. 2018).

The prevalence of transcripts of uncharacterized genes in the *in situ* transcriptomes (Figure 5B; Table S3) is particularly interesting. *C. parapsilosis* and other Candida species are rarely studied in a microbial community context, leaving gaps in understanding of genes required for organism-organism interactions. We suspect that some of the highly expressed genes are important for Candida interactions with bacteria and other community members. Thus, they represent important targets for future co-culture-based investigations.

## CONCLUSIONS

We applied genome-resolved metagenomics, metatranscriptomics, and metaproteomics to recover genomes for, and study the behavior of, *C. parapsilosis in situ*. We showed *C. parapsilosis* has a highly distinct transcriptomic profile *in situ* vs in culture. Further, the extreme variability in the *in situ* transcriptome data indicates the considerable effect the gut microbial community and human host may have on *C. parapsilosis* behavior and metabolism. Overall, these results demonstrate that *in situ* study of *C. parapsilosis* and other Candida species is not only possible but necessary for a more holistic understanding of their biology.

## METHODS

### Metagenomic sampling and sequencing

This study made use of previously published infant datasets: NIH1 (Brown et al. 2018), NIH2 (Brooks et al. 2017), NIH3 (Raveh-Sadka et al. 2015), NIH4 (Rahman et al. 2018), Sloan2 (Brooks et al. 2017), and SP_CRL (Sharon et al. 2013), as well as several new datasets including multiple timepoints from infant 06 and infant 74, and samples L2_023, S3_003, and S3_016.

For newly generated metagenomic sequencing from infant 06 and infant 74, total genomic DNA and total RNA were extracted from fecal samples using Qiagen's AllPrep PowerFecal DNA/RNA kit (Qiagen) and subsequently split into DNA and RNA portions. The aliquot used for metagenomic sample preparation was treated with RNase A. DNA quality and concentration were verified with Qubit (Thermofisher) and Fragment Analyzer (Agilent). Illumina libraries with an average insert size of 300 bps were constructed from purified genomic DNA using the Nextera XT kit (Illumina) and sequenced on Illumina's NovaSeq platform in a paired end 140 bp read configuration, resulting in at least 130 million paired end reads from each library.

NICU metagenomic sampling was described and published previously (Brooks et al. 2017). All samples were collected from the same NICU at UPMC Magee-Womens Hospital (Pittsburgh, PA). In order to generate enough DNA for metagenomic sequencing, DNA was collected from multiple sites in the NICU and combined into three separate pools for sequencing. Highly-touched surfaces included samples originating from the isolette handrail, isolette knobs, nurses hands, in-room phone, chair armrest, computer mouse, computer monitor, and computer keyboard. Sink samples included samples from the bottom of the sink basin and drain. Counters and floors consisted of the room floor and surface of the isolette. See previous publication for details (Brooks et al. 2017; Brooks et al. 2018).

### Eukaryotic genome binning and gene prediction

For each sample, sequencing reads were assembled independently with IDBA-UD (Peng et al. 2012). Additionally, for each infant, reads from every time point were concatenated together. A co-assembly was then performed on the pooled reads for each infant with IDBA-UD in order to assemble sequences from low abundance organisms. The Eukaryotic porton of each sample assembly was predicted with EukRep (West et al. 2018) and putative eukaryotic bins were generated by running CONCOCT (Alneberg et al. 2014) with default settings on the output of EukRep. To reduce computational load, resulting eukaryotic bins shorter than 2.5 mbp in length were not included in further analyses. GeneMark-ES (Ter-Hovhannisyan et al. 2008) and AUGUSTUS (Stanke et al. 2006) trained with BUSCO (Simão et al. 2015) were used to perform gene prediction on each bin using the MAKER (Cantarel et al. 2008) pipeline. In addition, a second homology-based gene prediction step was performed. Each bin was identified as either *C. parapsilosis* or *C. albicans* and reference gene sets from *C. parapsilosis CDC317* and *C. albicans SC5314* were used for homology evidence respectively in a second-pass gene prediction step with AUGUSTUS (Stanke et al. 2006), as implemented in MAKER (Cantarel et al. 2008).

### SNV calling and detection of SNV hotspots

In order to call variants in each Candida genome, reads from the sample in which a particular genome was binned from, or the publically available reads from SRA, were mapped back to the *de novo* assembled genome using Bowtie 2 (Langmead et al. 2012) with default parameters. The PicardTool (http://broadinstitute.github.io/picard/) functions “SortSam” and “MarkDuplicates” were used to sort the resulting sam file and remove duplicate reads. FreeBayes (Garrison et al. 2012) was used to perform variant calling with the options “--pooled-continuous −F 0.01 −C 1.” Variants were filtered downstream to include only those with support of at least 10% of total mapped reads in order to avoid false positives. SNV read counts were calculated using the “AO” and “RO” fields in the FreeBayes vcf output file.

SNV density was visualized across the CDC317 reference genome using a custom python script (https://github.com/patrickwest/c_parapsilosis_analysis). SNV hotspots were quantitatively defined with 5 kbp windows with a slide of 500 bp across the genome, flagging windows with a SNV density at least three standard deviations above the genomic average SNV density, and merging overlapping flagged windows. Genes located within SNV hotspots as well as overlapping SNV hotspots between strains were identified with intersectBed (Quinlan et al. 2010).

### Candida phylogenetics and population structure

For generation of a SNP tree for both *C. parapsilosis* and *C. albicans*, all publically available genomic sequencing reads for both species were downloaded from NCBI’s short read archive (SRA), including X isolate *C. parapsilosis* read sets and X *C. albicans* sets. SNVs were called for each isolate read set using the same pipeline used for metagenome-derived genomes, as described above. A SNP tree was generated for *C. parapsilosis* and *C. albicans* using SNPhylo (Lee et al. 2014) with settings ‘-r −M 0.5 −l 2’ and ‘-r −M 0.5 −l 0.8’ respectively and visualized using FigTree (https://github.com/rambaut/figtree/). For genomic average nucleotide identity (ANI) comparisons, X *C. parapsilosis* and X *C. albicans* reference genomes were downloaded from NCBI. Subsequently, dRep (Olm et al. 2017) in the ‘compare_wf’ setting was used to generate ANI comparisons for each genome. For inferring *C. parapsilosis* population structure, FreeBayes vcf files were converted to PLINK bed format with PLINK (Pucell et al. 2007) and used as input for ADMIXTURE (Alexander et al. 2011). The predicted number of ancestral populations, K, was selected using ADMIXTURE’s cross-validation procedure for values 1-8.

### Detection of copy number variation

Genomic copy number variation within the *C. parapsilosis* strains was searched for by mapping reads from the sample the genome was derived from to the *C. parapsilosis* CDC317 reference genome. Windowed coverage was then calculated across the genome in 100bp sliding windows using pipeCoverage (https://github.com/MrOlm/pipeCoverage) and visualized with Integrated Genomics Viewer (IGV) (Robinson et al. 2017). Copy numbers for multicopy regions were estimated by dividing the average coverage of the windows located within the multicopy region by the average genomic coverage.

### Transcriptomic sequencing and analysis

Total RNA was extracted from fecal samples using the AllPrep PowerFecal DNA/RNA kit (Qiagen) and subsequently treated with DNase. Purified RNA quality and concentration were measured using the Fragment Analyzer (Agilent). Illumina sequencing libraries were constructed with the ScriptSeq Complete Gold Kit (Illumina) without performing the rRNA removal step, resulting in library molecules with an average insert size of 150 bp. Sequencing was performed on Illumina's NextSeq platform in a paired end 75 bp configuration, resulting in an average of 54 million paired end reads per sample.

Transcriptomic reads from studies [ref][ref][ref] were downloaded from the SRA. Transcriptomic reads from each dataset were then mapped to *C. parapsilosis* reference strain CDC317 gene models with Kallisto (Bray et al. 2016) and transcript per million (TPM) values were used to compare expression levels across samples. Differentially expressed transcripts were identified using raw read counts with the R package DESeq2 (Love et al. 2014). Rlog transformation was applied to transcript read counts from each sample prior to generation of transcriptome PCAs. PCA plots were generated with DESeq2. Important features for separating C. parapsilosis *in situ* and culture as well as *E. faecalis* and *S. epidermidis* Candida-positive and Candida-negative samples were identified through the use of a sparse Partial Least Squares Discriminant Analysis (sPLS-DA) as implemented in the MixOmics package (Rohart et al. 2017) on rlog transformed transcript read counts. MixOmics cross-validation (tune.splsda) was used with settings fold = 3 and nrepeat = 50 to estimate the optimal number of components (features) for separating each pair of sample types.

Genes were annotated with KEGG KOs and PFAM domains using HMMER with KOfam (Aramaki et al. (2019) and Pfam-A (El-Gebali et al. 2019) HMM databases. Subsets of genes of interest (described in results) were then searched for significantly enriched KEGG families or PFAM domains with a hypergeometric distribution test as part of the R ‘stats’ package (R Core Team, 2013).

### Generation of Proteomic Datasets

Lysates were prepared from ~50mg of fecal material by bead beating in SDS buffer (4% SDS, 100 mM Tris-HCl, pH 8.0) using 0.15-mm diameter zirconium oxide beads. Cell debris was cleared by centrifugation (21,000 x g for 10 min). Pre-cleared protein lysates were adjusted to 25mM dithiothreitol and incubated at 85°C for 10 min to further denature proteins and to reduce disulfide bonds. Cysteine residues were alkylated with 75 mM iodoacetamide, followed by a 20-minute incubation at room temperature in the dark. After incubation, proteins were isolated by chloroform-methanol extraction. Protein pellets were washed with methanol, air-dried, and resolubilized in 4% sodium deoxycholate (SDC) in 100 mM ammonium bicarbonate (ABC) buffer, pH 8.0. Protein samples were quantified by BCA assay (Pierce) and transferred to a 10 kDa MWCO spin filter (Vivaspin 500; Sartorius) before centrifugation at 12,000 x g to collect denatured and reduced proteins atop the filter membrane. The concentrated proteins were washed with 100 mM ABC (2x the initial sample volume) followed by centrifugation. Proteins were resuspended in a 1x volume of ABC before proteolytic digestion. Protein samples were digested in situ using sequencing-grade trypsin (G-Biosciences) at a 1:75 (wt/wt) ratio and incubated at 37°C overnight. Samples were diluted with a 1x volume of 100 mM ABC, supplied with another 1:75 (wt/wt) aliquot of trypsin, and incubated at 37°C for an additional 3 hours. Tryptic peptides were then spin-filtered through the MWCO membrane and acidified to 1% formic acid to precipitate the residual SDC. The SDC precipitate was removed from the peptide solution with water-saturated ethyl acetate extraction. Samples were concentrated via SpeedVac (Thermo Fisher), and peptides were quantified by BCA assay (Pierce) before LC-MS/MS analysis.

12ug of each peptide sample was analyzed by automated 2D LC-MS/MS using a Vanquish UHPLC with autosampler plumbed directly in-line with a Q Exactive Plus mass spectrometer (Thermo Scientific). A 100 μm inner diameter (ID) triphasic back column [RP-SCX-RP; reversed-phase (5 μm Kinetex C18) and strong-cation exchange (5 μm Luna SCX) chromatographic resins; Phenomenex] was coupled to an in-house pulled, 75 μm ID nanospray emitter packed with 30 cm Kinetex C18 resin. Peptides were autoloaded, desalted, separated, and analyzed across four successive salt cuts of ammonium acetate (35, 50, 100, and 500 mM), each followed by a 105-minute organic gradient. Mass spectra were acquired in a data-dependent mode with the following parameters: a mass range of 400 to 1,500 m/z; MS and MS/MS resolution of 35K and 17.5K, respectively; isolation window = 2.2 m/z with a 0.5m/z isolation offset; unassigned charges and charge states of +1, + 5, +6, +7 and +8 were excluded; dynamic exclusion was enabled with a mass exclusion window of 10 ppm and an exclusion duration of 45seconds.

MS/MS spectra were searched against custom-built databases composed of the concatenated sequenced metagenome derived predicted proteomes from all time-points, the human reference proteome from UniProt, common protein contaminants, and reversed-decoy sequences using Proteome Discover 2.2 (Thermo Scientific), employing the CharmeRT workflow (Dorfer et al., 2018). Peptide spectrum matches (PSMs) were required to be fully tryptic with two miscleavages, a static modification of 57.0214 Da on cysteine (carbamidomethylated) residues, and a dynamic modification of 15.9949 Da on methionine (oxidized) residues. False-discovery rates (FDRs), as assessed by matches to decoy sequences, were initially controlled at 1% at the peptide level. To alleviate the ambiguity associated with shared peptides, proteins were clustered into protein groups by 100% identity for microbial proteins and 90% amino acid sequence identity for human proteins using USEARCH (Edgar et al., 2010). FDR-controlled peptides were then quantified according to the chromatographic area under the curve (AUC) and mapped to their respective proteins. Peptide intensities were summed to estimate protein-level abundance based on peptides that uniquely mapped to one protein group. Protein abundance distributions were then normalized across samples using InfernoRDN (Polpitiya et al, 2008), and missing values were imputed to simulate the mass spectrometer’s limit of detection using Perseus (Tyanova et al., 2016).

## DECLARATIONS

### Ethics approval and consent to participate

This study was reviewed and approved by the University of Pittsburgh Institutional Review Board (IRB PRO12100487 and PRO10090089).

### Consent for publication

Not applicable

### Availability of data and materials

The datasets supporting the conclusions of this article are available in the NCBI BioProeject repository, PRJNA471744 https://www.ncbi.nlm.nih.gov/bioproject/?term=PRJNA471744, the Short Read Archive (SRA) SRR5420274 to SRR5420297, https://www.ncbi.nlm.nih.gov/sra/?term=SRR5420274, and https://github.com/patrickwest/c_parapsilosis_analysis. Several new datasets are still in the process of being uploaded to NCBI and SRA and will have their accession numbers added before publication. Further, Candida genomes are available via https://ggkbase.berkeley.edu/project_groups/candida_genomes (sign in as a user is required).

### Competing interests

The authors declare that they have no competing interests

### Funding

This research was supported by the National Institutes of Health (NIH) under award RAI092531A, the Alfred P. Sloan Foundation under grant APSF-2012-10-05, and National Science Foundation Graduate Research Fellowships to P.W. under Grant No. DGE 1106400. This work used the Vincent J. Coates Genomics Sequencing Laboratory at UC Berkeley, supported by NIH S10 OD018174 Instrumentation Grant.

## Acknowledgements

We thank Spencer Diamond and Alex Crits-Christoph for helpful discussions and advice on statistical methods as well as Michelle Tan, Rene Sit, and Norma Neff from Chan Zuckerberg Biohub Genomics Platform for providing sequencing resources.

## FIGURE CAPTIONS

Figure S1

Candida parapsilosis heterozygosity and ANI comparisons display low C. parapsilosis heterozygosity, even in a metagenome context, and a possible hospital associated population. (A) *C. parapsilosis* heterozygosity for various strains as measured by heterozygous variants per kb.(B) ANI comparisons for each *C. parapsilosis* strain. Comparisons made using dRep with the ‘compare_wf’ setting. If multiple genomes were assembled for the same strain from different time points, all assembled genomes were included in this analysis.

Figure S2

Phylogenetic and ANI comparisons of C. albicans genomes show no clear hospital associated population. (A) ANI comparisons for each C. albicans strain. All publically available C. albicans genomes from NCBI were included. Genomes assembled in this study are highlighted in red. If multiple genomes were assembled for the same strain from different time points, all assembled genomes were included in this analysis. (B) A phylogenetic tree of *C. albicans* strains constructed from concatenated SNVs. Strains with genomes assembled in this study are highlighted in red. All publically available C. albicans genome read sets on NCBI are included in this analysis.

Figure S3

Population structure analysis of C. parapsilosis strain genomes performed with ADMIXTURE (Alexander et al. 2011) reveals possible admixture in strain N3_182. Each color represents a different ancestral population (N=6) and the proportion of a color in each strain represents how much of the variation in its genome is attributed to that particular ancestral population. All strains except N3_182 are predicted to have a separate ancestral population.

Figure S4

Whole genome SNV density plots for each *C. parapsilosis* strain shows SNV hotspots regardless of reference genome used for mapping. Strain names on the far left represent the genome used for read mapping, whereas strain names on right represent the read set used for mapping. Scaffolds are separated with dashed lines. Total bar height represents total SNV density and homozygous SNV proportion is labeled in red whereas heterozygous is black.

Figure S5

sPLS-DA important feature selection. (A) Separation of sample categories (*in situ* vs culture and Candida+ vs Candida-) based on the selected number features and visualized using the first two components of the sPLS-DA. (B) Visualization of the correlation of selected features. Features projected in the same direction are correlated, with greater distance from the origin depicting a stronger correlation.

Figure S6

Expression of RTA3 over time in infant 06 for strain C1_006 shows no clear response to fluconazole treatment at the time points measured. Fluconazole treatment was administered to infant 06 at DOL 19 and caspofungin treatment at DOL 26.

Figure S7

C. parapsilosis *in situ* transcriptomes show more variance than that observed between C. albicans white and opaque phenotypes. Samples are hierarchically clustered, with top bars reflecting how similar samples are to one another. Y axis represents only transcripts differentially expressed between white and opaque phenotypes identified in Tuch et al. 2010. C. parapsilosis orthologs for each C. albicans transcript were identified with orthofinder.

