## Supplementary material for "Genetic and behavioral adaptation of *Candida parapsilosis* to the microbiome of hospitalized infants revealed by *in situ* genomics, transcriptomics and proteomics": Figure S1

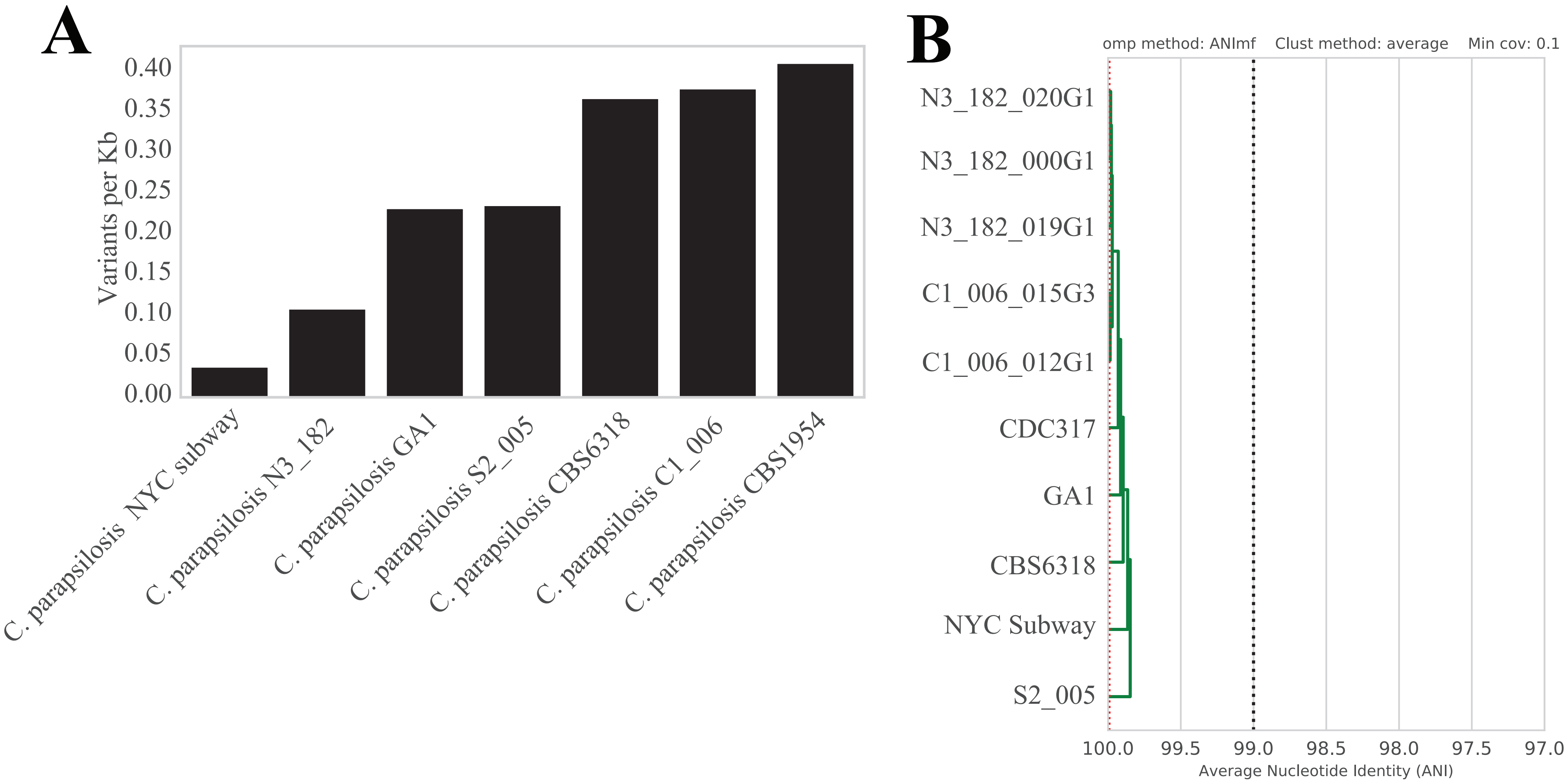


Figure S1: **Candida parapsilosis heterozygosity and ANI comparisons display low C. parapsilosis heterozygosity, even in a metagenome context, and a possible hospital associated population.** (A) C. parapsilosis heterozygosity for various strains as measured by heterozygous variants per kb.(B) ANI comparisons for each C. parapsilosis strain. Comparisons made using dRep with the ‘compare_wf’ setting. If multiple genomes were assembled for the same strain from different time points, all assembled genomes were included in this analysis.
