## Supplementary material for "Genetic and behavioral adaptation of *Candida parapsilosis* to the microbiome of hospitalized infants revealed by *in situ* genomics, transcriptomics and proteomics": Figure S2

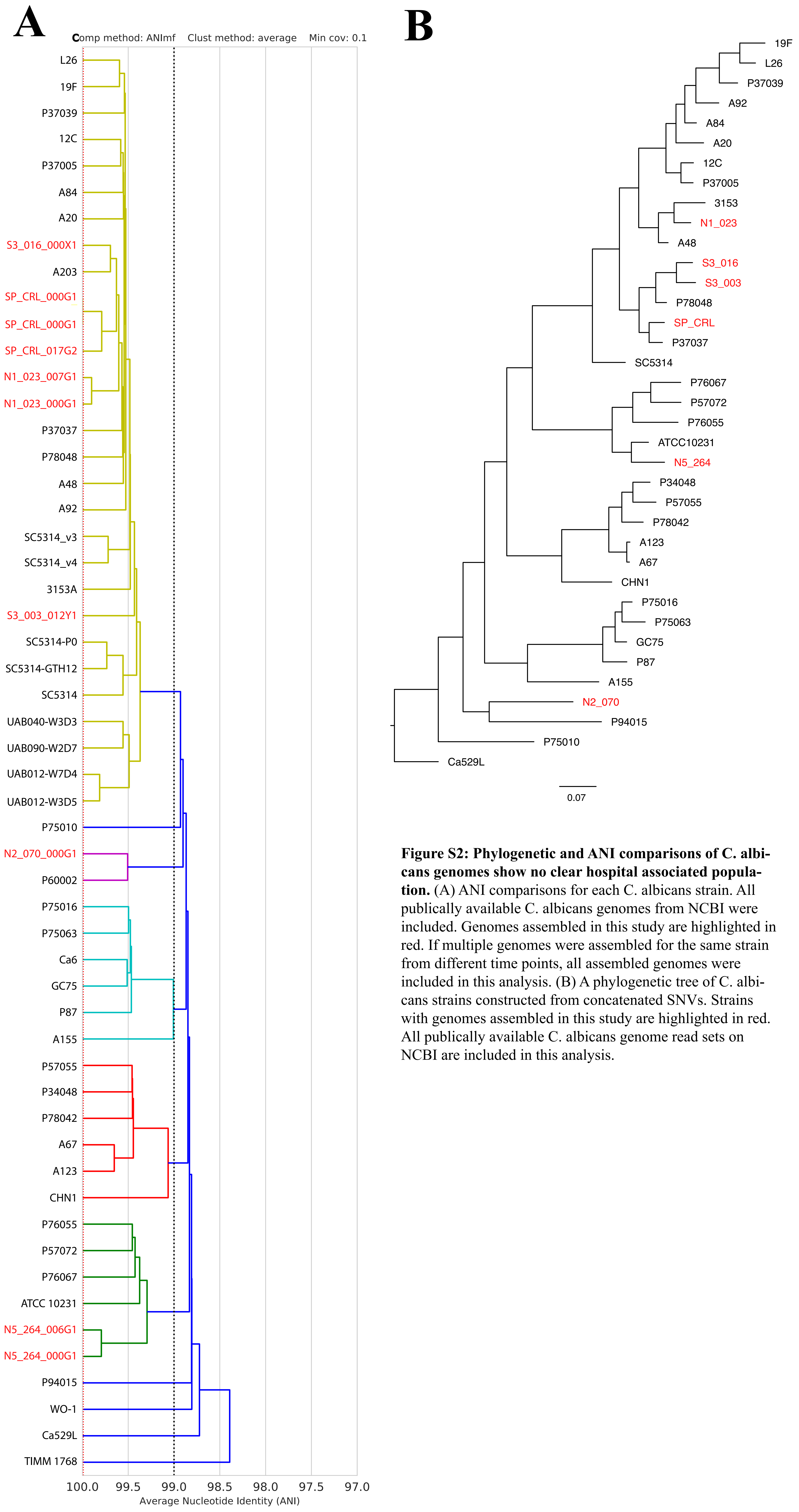


**Figure S2: Phylogenetic and ANI comparisons of C. albicans genomes show no clear hospital associated population.** (A) ANI comparisons for each C. albicans strain. All publically available C. albicans genomes from NCBI were included. Genomes assembled in this study are highlighted in red. If multiple genomes were assembled for the same strain from different time points, all assembled genomes were included in this analysis. (B) A phylogenetic tree of C. albicans strains constructed from concatenated SNVs. Strains with genomes assembled in this study are highlighted in red. All publically available C. albicans genome read sets on NCBI are included in this analysis.
