## Supplementary material for "Genetic and behavioral adaptation of *Candida parapsilosis* to the microbiome of hospitalized infants revealed by *in situ* genomics, transcriptomics and proteomics": Figure S3

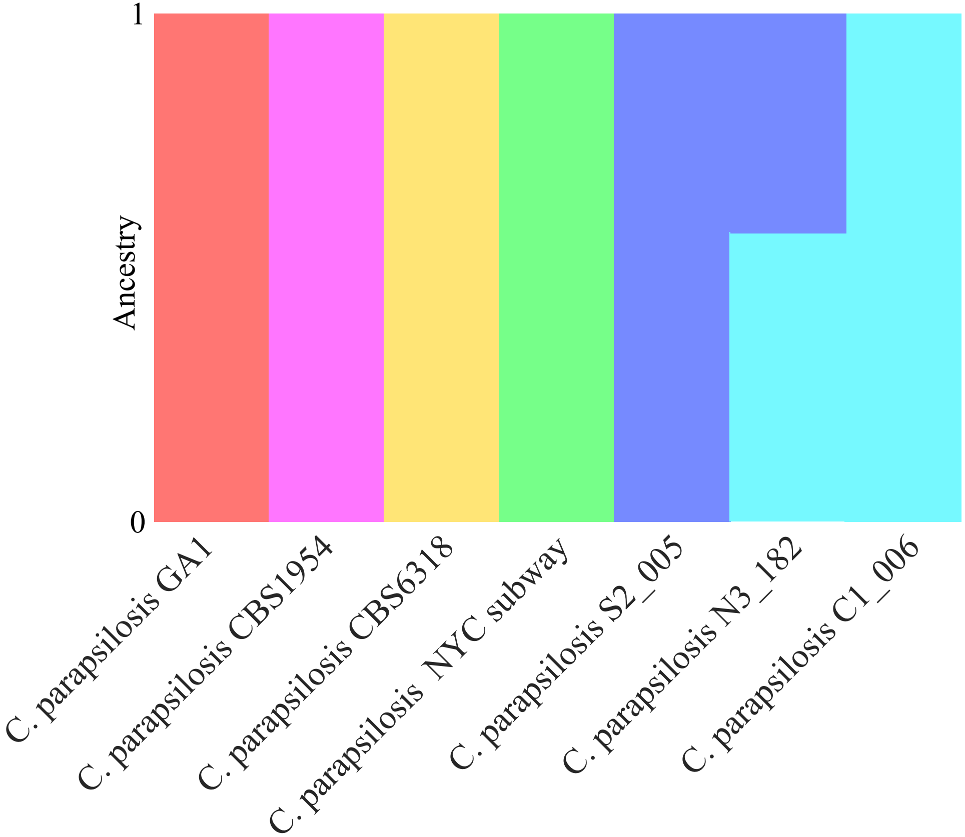


**Figure S3: Population structure analysis of C. parapsilosis strain genomes performed with ADMIXTURE (Alexander et al. 2011) reveals possible admixture in strain N3_182.** Each color represents a different ancestral population (N=6) and the proportion of a color in each strain represents how much of the variation in its genome is attributed to that particular ancestral population. All strains except N3_182 are predicted to have a separate ancestral population.
