## Supplementary material for "Genetic and behavioral adaptation of *Candida parapsilosis* to the microbiome of hospitalized infants revealed by *in situ* genomics, transcriptomics and proteomics": Figure S4

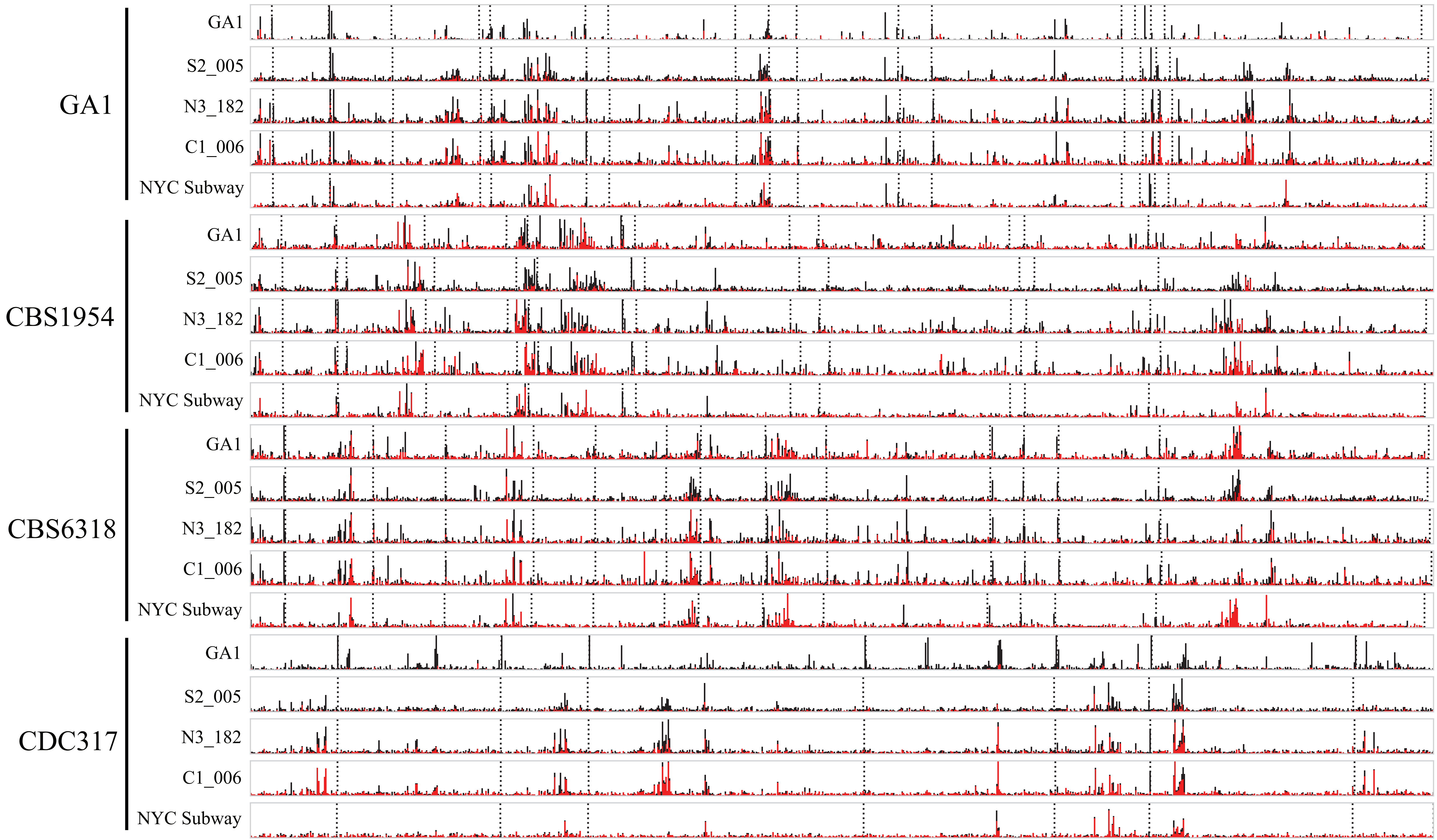


**Figure S4: Whole genome SNV density plots for each C. parapsilosis strain shows SNV hotspots regardless of reference genome used for mapping.** Strain names on the far left represent the genome used for read mapping, whereas strain names on right represent the read set used for mapping. Scaffolds are separated with dashed lines. Total bar height represents total SNV density and homozygous SNV proportion is labeled in red whereas heterozygous is black.
