## Supplementary material for "Genetic and behavioral adaptation of *Candida parapsilosis* to the microbiome of hospitalized infants revealed by *in situ* genomics, transcriptomics and proteomics": Figure S5

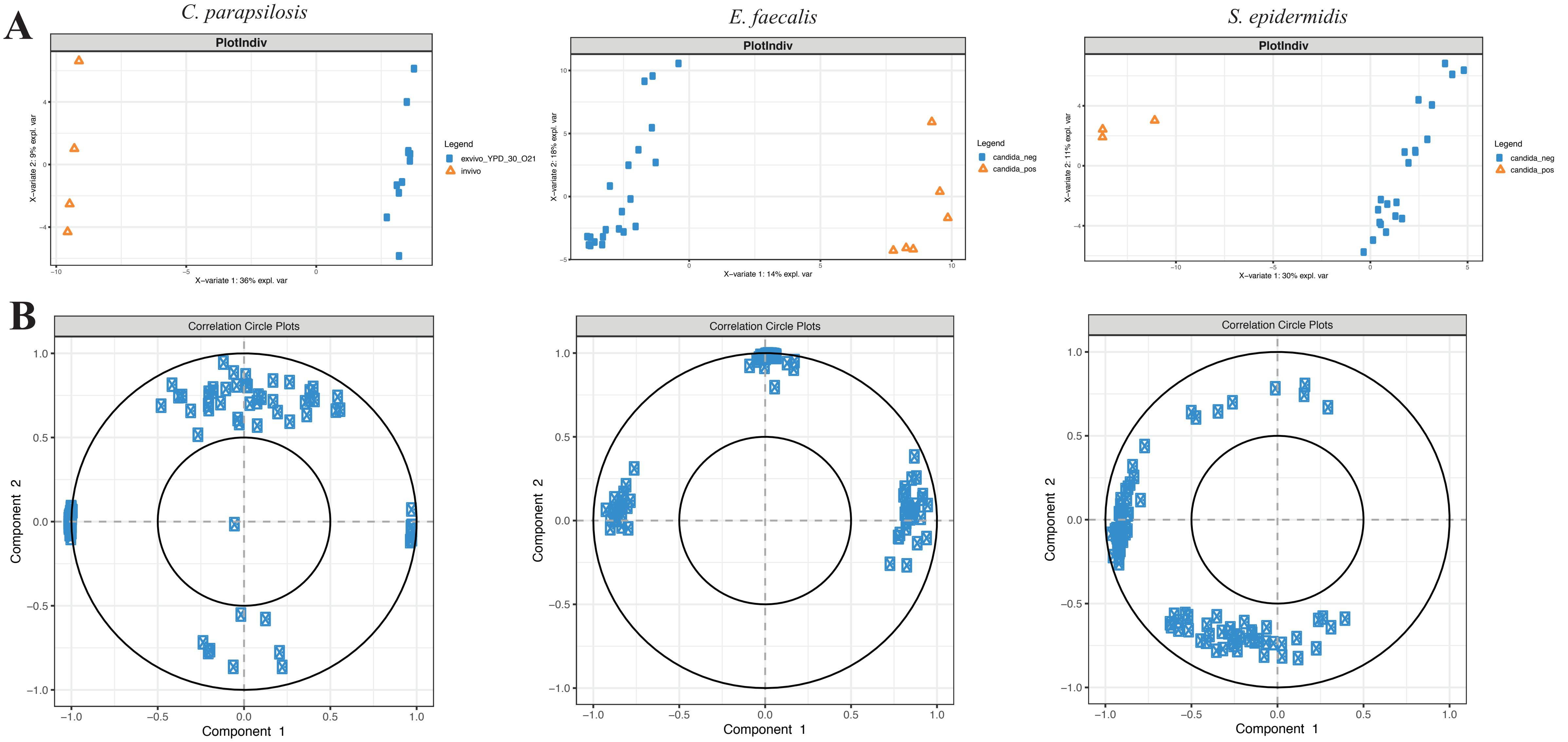


**Figure S5: sPLS-DA important feature selection.** (A) Separation of sample categories (in situ vs culture and Candida+ vs Candida-) based on the selected number features and visualized using the first two components of the sPLS-DA. (B) Visualization of the correlation of selected features. Features projected in the same direction are correlated, with greater distance from the origin depicting a stronger correlation.
