## Supplementary material for "Genetic and behavioral adaptation of *Candida parapsilosis* to the microbiome of hospitalized infants revealed by *in situ* genomics, transcriptomics and proteomics": Figure S6

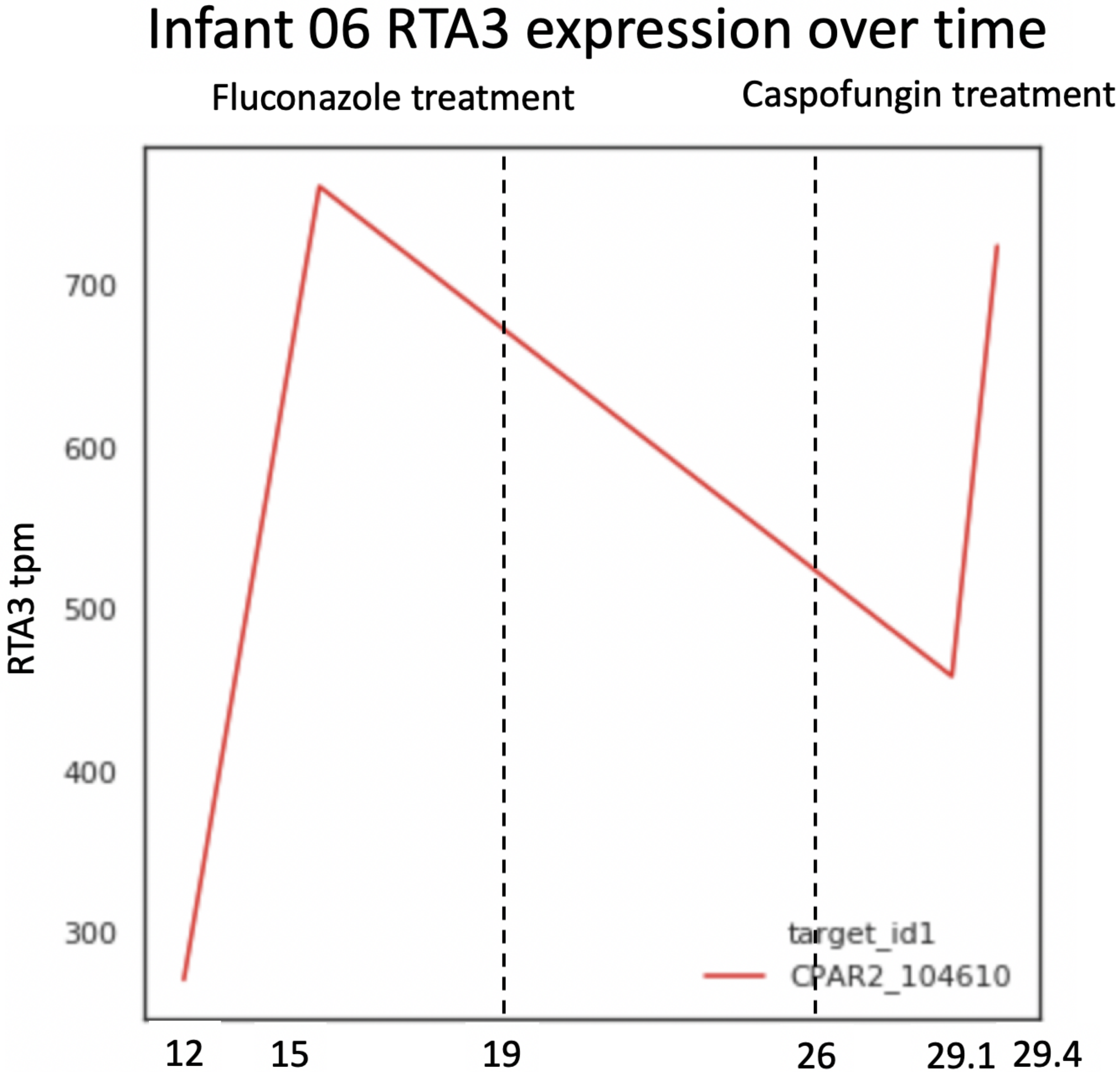


**Figure S6: Expression of RTA3 over time in infant 06 for strain C1_006 shows no clear response to fluconazole treatment at the time points measured.** Fluconazole treatment was administered to infant 06 at DOL 19 and caspofungin treatment at DOL 26.
