## Supplementary material for "Genetic and behavioral adaptation of *Candida parapsilosis* to the microbiome of hospitalized infants revealed by *in situ* genomics, transcriptomics and proteomics": Figure S7

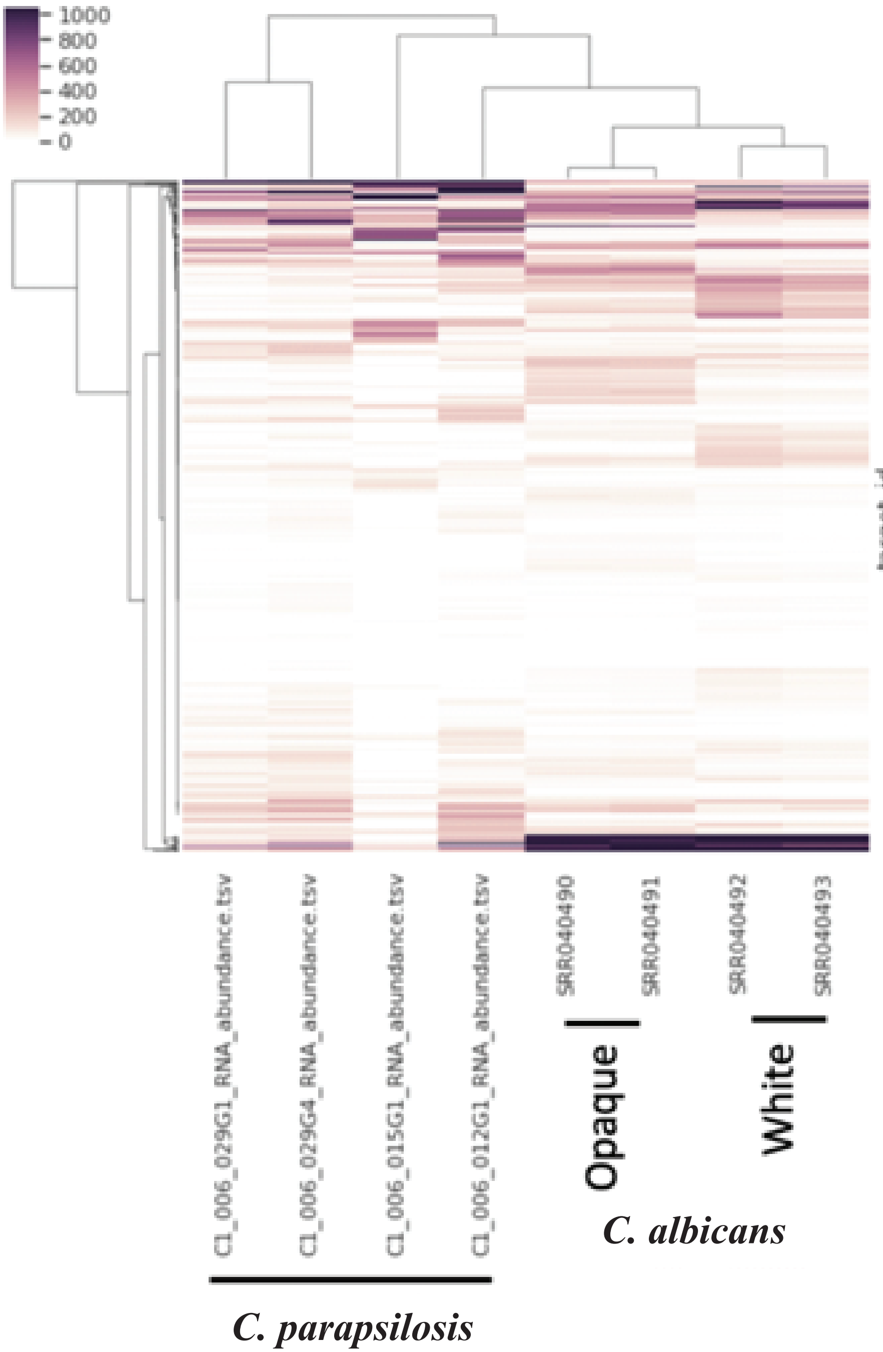


**Figure S7: C. parapsilosis in situ transcriptomes show more variance than that observed between C. albicans white and opaque phenotypes.** Samples are hierarchically clustered, with top bars reflecting how similar samples are to one another. Y axis represents only transcripts differentially expressed between white and opaque phenotypes identified in Tuch et al. 2010. C. parapsilosis orthologs for each C. albicans transcript were identified with orthofinder.
